# Foregut organ progenitors and their niche display distinct viscoelastic properties *in vivo* during early morphogenesis stages

**DOI:** 10.1101/2021.05.31.446410

**Authors:** Aliaksandr Dzementsei, Younes F. Barooji, Elke A. Ober, Lene B. Oddershede

**Author notes:** These authors contributed equally to this work (Aliaksandr Dzementsei, Younes F. Barooji). Corresponding authors’.

## Abstract

Material properties of living matter play an important role for biological function and development. Yet, quantification of material properties of internal organs *in vivo*, without causing physiological damage, remains challenging. Here, we present a non-invasive approach based on modified optical tweezers for quantifying sub-cellular material properties deep inside living zebrafish. Material properties of cells within the gut region of living zebrafish are quantified as deep as 150 *μ*m into the biological tissue. The measurements demonstrate differential mechanical properties of the developing foregut organs progenitors: Gut progenitors are more elastic than any of the neighboring cell populations at the time when the developing organs undergo substantial displacements during morphogenesis. The higher elasticity of gut progenitors correlates with an increased cellular concentration of microtubules. The results infer a role of material properties during morphogenesis and the approach paves the way for quantitative material investigations *in vivo* of embryos, explants, or organoids.

## Introduction

Material properties are crucial for action-reaction mechanisms and are closely linked to motion, also within living organisms. Viscoelasticity, short for the combined viscous and elastic properties of a material, determines the fluidity and stiffness of a material and the way it responds to internally and externally generated forces(1). Furthermore, viscoelasticity has been shown to correlate, e.g., with the invasiveness of cancer(2) and with the differentiation of stem cells(3, 4). The ability of cells to move or maintain or change their shape is in part regulated by their reaction to external forces depending on the cells’ material properties (5). However, the material properties of cells, tissues or organs are largely unknown and a quantification of these is the first step in understanding how material properties may contribute to organ and overall embryo morphogenesis.

The contribution of mechanical forces and biophysical material properties, such as viscoelasticity, remains poorly understood, predominantly due to a lack of tools to measure and quantify material properties within complex biological systems *in vivo*. Several assays have been developed to analyze micro-rheological properties of cells cultured *in vitro* or of surface embryonic tissues, such as atomic force microscopy, micro-aspiration or optical tweezers (6). These assays can be used to quantify the viscous and elastic properties of living matter on a variety of time and length scales down to micro-seconds and the sub-cellular level (7–9). However, *in vivo* investigation of material properties of cells and tissues forming internal organs remains difficult due to the challenge of performing accurate quantitative measurements, especially deep within a living organism without causing physiological damage to the investigated organism.

Optical tweezers are widely used *in vitro* to investigate protein folding (10), and also to quantify material properties of isolated cells (11). In its simplest form, an optical trap is formed by tightly focusing a laser beam, whereby objects with a refractive index higher than the surroundings are drawn along the intensity gradient towards the focus of the beam by a harmonic force (12). The near-infrared (NIR) biological transparency window allows NIR lasers to penetrate deeply into biological tissues and if NIR-based optical traps are operated at sufficiently low laser powers, they cause no observable physiological damage (13). Recently, there have been reports of successful optical manipulation of particles (14, 15), red blood cells (16) and otoliths (17) in living zebrafish embryos. However, most of these studies were qualitative and the deepest reported trapping site was about 50 *μ*m inside living zebrafish (14). In the early *Drosophila* embryo, optical tweezers were used to probe the mechanics of cell contacts close to the surface of the organism by observing the equilibration of the interface following a deformation (18). A quantitative approach combining active oscillations and passive measurements was developed based on optical tweezers to determine viscous and elastic parameters inside living cultured cells (19). A similar method was used to determine the ratio between the viscous and elastic moduli within cells close to the surface of early zebrafish embryos at 5-7 hours post fertilization (hpf) (15). Other approaches to infer viscoelasticity inside living organisms are based on video recordings of either magnetically responsive ferrofluid microdroplets in zebrafish tailbuds (20) or thermal fluctuations of particles in the syncytium of the early *Drosophila* embryos (21). These methods were used to determine viscoelastic properties relatively close to the surface of the embryo and cannot directly be applied to investigate internal tissues and organs *in vivo* due to light scattering and poor resolution deep inside biological tissues.

Here, we measure material properties at sub-cellular resolution in deep tissues using laser tracking and photodiode detection of the thermal fluctuations of optically trapped nanoparticles. To investigate whether material properties differ between cell types undergoing complex morphogenetic movements, we focused on liver and foregut morphogenesis in zebrafish at the point in time when organ asymmetry is established by rearrangement of four cell populations next to the extraembryonic yolk (22). The liver and gut progenitor populations arise from the foregut endoderm. After their specification, liver progenitors form an asymmetric organ primordium, the liver bud, by directional cell migration to the left of the midline (Figure 1A) (23). In a parallel process, the entire foregut undergoes leftward looping, which is suggested to result from a passive displacement caused by asymmetric movements of the neighboring left and right lateral plate mesoderm (LPM) (24). These morphogenetic movements ensure efficient arrangement of the organs within the body cavity, which is crucial for proper physiological function. The mechanical properties of these developing organs have not yet been reported, probably because they are located 120-150 *μ*m deep inside the embryo (Figure 1B-C)(22, 25). We demonstrate consistent and significant differences in cellular viscoelasticity between the analyzed progenitor populations, with the gut progenitors being the most elastic in comparison to neighboring rearranging tissues. The time-scales probed in the current investigation are those relevant for cytoskeletal biopolymer dynamics inside living cells. Our data show that the observed biomechanical differences correlate with different concentrations of microtubules, hence, material properties may influence morphogenetic tissue movements during organ formation and embryonic development.

**Figure 1.**
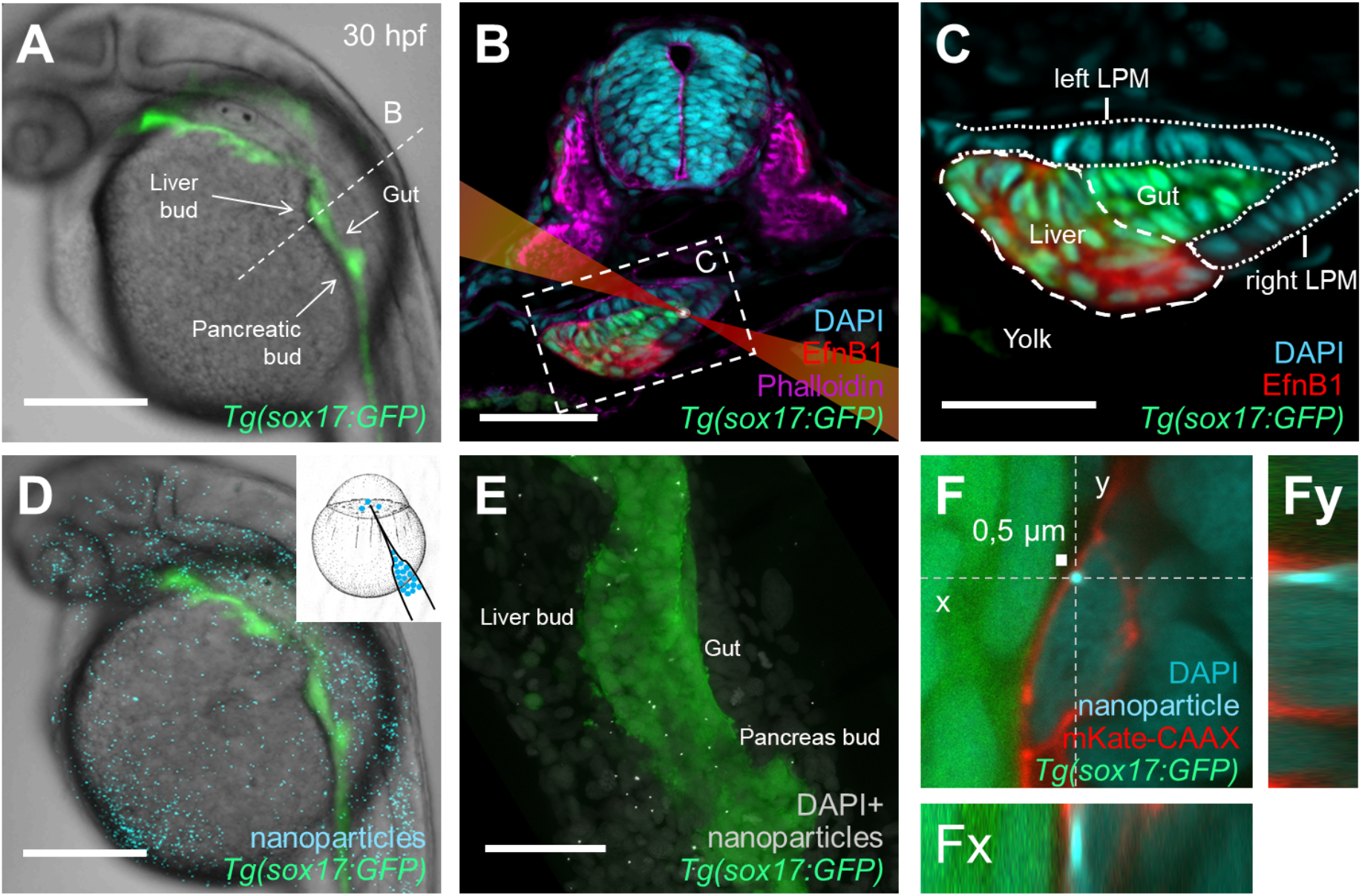
Injected nanoparticles distribute throughout the embryo, including the foregut region in 28-30 hpf zebrafish embryos. (A) Transgenic *tg(−0.5 sox17:GFP)^zf99^* visualizes the endoderm in living zebrafish embryo, including the foregut organ primordia. (B) Transverse section through the foregut region (dashed line in A) with schematized laser beam path (red) and bead (white). The foregut endoderm expresses transgenic *sox17:GFP* (green) and the liver progenitors EphrinB1 (red); Dapi marks the nuclei (cyan); Phalloidin marks F-actin (magenta). (C) Magnification of the foregut region shown in B (white dashed rectangle). (D) 0.5 *μ*m polystyrene particles (cyan) microinjected at the one-cell stage distribute throughout the embryo, including the endoderm (green), without causing apparent morphological defects. (E) Projection of confocal stack showing distribution of nanoparticles (white) in the foregut region, GFP marks the endoderm. (F) Confocal image of a representative nanoparticle (light blue) located in the cytoplasm between nucleus (darker blue) and plasma membrane (red) in a fixed embryo. (Fy) and (Fx) show orthogonal views in the ‘yz’ plane (dashed line ‘y’ in F) and ‘xz’ plane (dashed line ‘x’ in F), respectively. Scale bars A, D: 250 *μ*m; B, E: 50 *μ*m; C: 30 *μ*m; F: 0,5 *μ*m (scale bar next to bead).

## RESULTS

### Micro-injected nanoparticles can be optically trapped deep within living zebrafish embryos

Viscoelastic properties of a matrix, such as the intracellular environment, can be mapped by optical tweezers based tracking of thermal fluctuations of particles within the matrix (7, 8, 26). Intracellular organelles like lipid granules can serve as trackable endogenous particles and have successfully been used for probing single cell systems (8, 27). However, *in vivo* tracking of lipid granules is difficult in the majority of deep tissues as their signal-to-noise ratio is not high and it decreases with increasing penetration depth due to increasing spherical aberration.

In our study, we assessed the suitability of 0.2*μ*m gold and 0.5, 0.8 and 1 *μ*m fluorescently-labelled polystyrene nanoparticles (all numbers given are the diameters of the particles) as tracers and introduced them by microinjection into 1-cell stage zebrafish embryos. During embryonic development, the 0.5 mm polystyrene beads distributed throughout the embryo without causing any apparent morphological defects at 30 hpf (Figure 1D-E) or 5 days post fertilization, as determined by bright-field stereomicroscopy (data not shown). In the foregut region, the majority of these nanoparticles were located intracellularly with typically only one bead per cell positioned between the nucleus and the plasma membrane (Figure 1F). In contrast, gold nanoparticles, as well as 0.8 and 1 *μ*m polystyrene beads were rarely or not detected in the foregut region. The latter could be due to the large particle size compared to the relatively small progenitor cells in the foregut region.

To minimize scattering of both NIR and visible light during the *in vivo* measurements, the injected embryos at the stage of liver bud formation were dorsolaterally embedded in agarose and placed on the stage of an inverted confocal microscope with the liver bud facing the microscope objective; see orientation of zebrafish with respect to the trapping laser path in Figure 1B and 2A (inset). Injected nanoparticles could be optically trapped in diverse tissues, including somites (muscle cells), spinal cord (neurons) and the foregut region (endoderm and lateral plate mesoderm), by a tightly focused NIR laser beam (1064 nm) implemented in a confocal microscope (Figure 2A)(28), which allows for simultaneous optical trapping and confocal visualization of the fluorescent nanoparticles. To demonstrate optical trapping of nanoparticles in the foregut region, the embedded zebrafish embryo was moved at low speed by a piezo stage relative to the trapping laser. When an internalized particle co-localized with the optical trap, the particle became visibly trapped and remained trapped for several seconds in spite of the continued movement of the embryo (Supplementary Video). Likewise, the difference in trajectories of nanoparticles in the zebrafish foregut in trapped or freely diffusing states indicates that the NIR optical tweezers can trap particles at considerable depth in living embryos (inset of Figure 2B).

**Figure 2.**
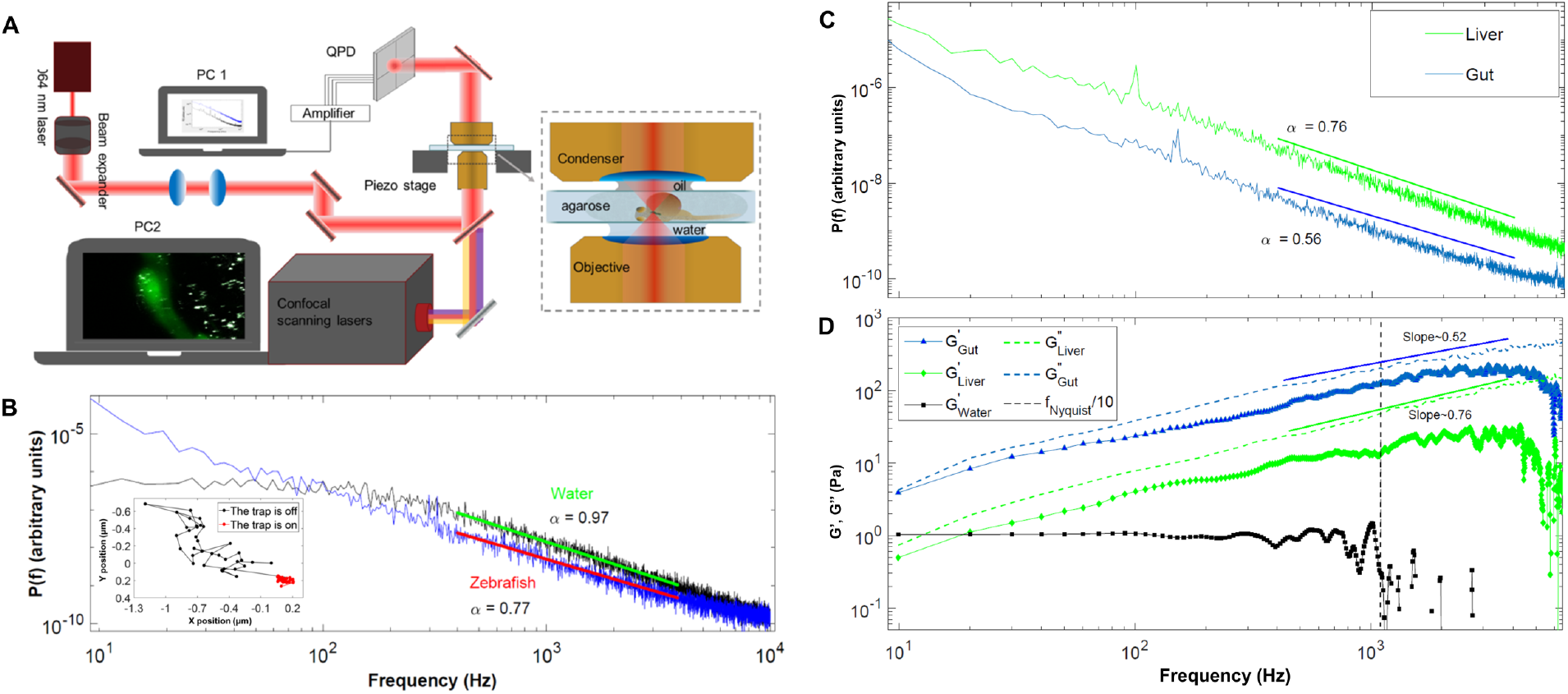
Forward scattered light from optically trapped nanoparticles in living zebrafish can be used to infer cellular viscoelasticy. (A) Schematic of the setup for laser tracking of nanoparticles *in vivo*: the laser beam (1064 nm, red line) is focused in the embryo (inset) and the forward-scattered light is collected by a condenser and imaged onto a quadrant photodiode (QPD). (B) Examples of positional power spectra of optically trapped particles in water (black, *α*=0.97) or in a zebrafish embryo (blue, *α*=0.77). The straight lines show fits of equation (1) to data. The inset shows the trajectory of a particle in the foregut region of a living zebrafish embryo where the trap is first on (red) or then off (black). (C) Power spectra of optically trapped nanoparticles in liver (green) or gut progenitors (blue). Full lines show linear fits to the double logarithmic plot in the frequency interval 400 Hz < f < 4000 Hz, yielding scaling exponents of 0.56±0.08 (gut) and 0.76±0.06 (liver), respectively. (D) Symbols (blue and green) provide the storage moduli, G’, of the gut and liver, same colors as in (C) and (D). The corresponding loss moduli, G”, are shown with dashed lines for comparison of the amplitude. G’’ dominates G’ over the entire frequency interval for both tissue types. The full lines show fits to the loss moduli data in the same region as fitted in the power spectra (C), returning scaling exponents of 0.52±0.09 (gut) and 0.76±0.05 (liver). Black squares denote the storage modulus as a function of frequency for a trapped bead in water. Water is a purely viscous liquid with a storage modulus of zero. The value of 1 Pa in this graph (black squares) denotes the constant storage modulus of the optical trapping potential, which can be reliably calculated for *f < f_Nyquist_/10* (dashed vertical line). The data shown in (C)-(D) is an average of 5 experiments for each cell type.

### Thermal fluctuations of optically trapped nanoparticles enable quantification of cellular viscoelasticity in internal tissues

Measurements of the viscoelastic properties of the foregut region were based on monitoring thermal fluctuations of microscopic tracers (8, 26, 29). To apply this method deep inside the living zebrafish embryo, we co-localized the tightly focused laser, the optical trap, and a nanoparticle in the desired tissue in three dimensions and recorded the positions of the particle’s thermal fluctuations by a quadrant photodiode (QPD) without moving the chamber (Figure 2A). The nanoparticle has a focusing effect on the laser light, therefore the signal detected by the QPD significantly increases when a nanoparticle was in the center of the trap compared to an empty trap within the embryo (Supporting Figure S1). To visually localize the foregut region in the embryo, we utilized transgenic *sox17:GFP* embryos with GFP expression highlighting the endoderm, including the liver and gut progenitors (Figure 1A). *In vivo* micro-rheological measurements were performed using several nanoparticles per embryo. The measurements took place at the time of asymmetric liver bud formation, around 30 hpf (Figure 1A-C). Subsequently, the embryo was fixed and the foregut region was imaged with improved resolution by confocal microscopy to determine the location of each of the tracked particles (Figure 1F). Particles occasionally observed in the extra-cellular space, as well as particles that could not be clearly assigned to a specific tissue, were excluded from the analysis.

The power spectrum, *P_x_*(*f*), of a tracer particle in a matrix can be calculated from its positional time series (Figure 2C). The power spectrum of a particle in a purely viscous medium, such as water, is well fitted by a Lorentzian function(30, 31), consistent with the behavior shown in Figure 2B with a flat region at low frequencies, reflecting motion where the tracer particle feels the restoring potential of the optical trap, and a scaling region at higher frequencies; the frequency separating these two regimes is denoted the corner frequency, *f_c_*. The scaling region with frequencies above *f_c_* carries information about the material properties of the matrix, which includes viscous and elastic properties (8, 26, 32–34). At frequencies above the corner frequency, the power spectra, *P_x_*(*f*) (Figure 2C), scale with frequency, *f*:

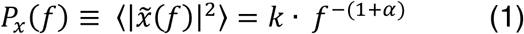

The scaling exponent, *a*, characterizes the scaling of the mean squared displacement (MSD) of the nanoparticle as a function of time, *t*: MSD = 〈|*x*(*t*)|^2^〉 ∝ *t^α^* (8, 32, 33). The scaling exponent provides information about the motion of the tracer: *α* = 0 signifies complete confinement, 0 < *α* < 1 indicates subdiffusion in a viscoelastic medium, and *α* = 1 is a sign of Brownian motion in a purely viscous medium, such as water. When comparing values of scaling exponents, a, in the subdiffusive regime (0<a<1), and if probing at time-scales where non-equilibrium processes are negligible, a higher value of a indicates a more viscous environment and a lower a value indicates a more elastic environment. The constant *k* from equation (1) contains information about the signal to noise ratio (as visible in Supporting Figure S1) and does not enter the calculation of *a*.

In the current study we focus on the intracellular material properties of organ progenitor cells and not on non-equilibrium processes such as cell motility. Therefore, to obtain a, we fitted the power spectra in a frequency an interval between 400-4000 Hz (corresponding to the 0.25-2.5 ms regime) where non-equilibrium processes are considered negligible and which is the frequency regime relevant for cytoskeletal polymer dynamics (21, 35, 36). This fitting interval is within the scaling region which is well above *f_c_* and below the 3dB cut-off frequency, *f_3dB_*, of the quadrant photodiode (37). A typical power spectrum resulting from optical trapping of a microinjected nanoparticle at a depth of about 150 μm in a living zebrafish, as well as a power spectrum obtained by trapping a similar nanoparticle in water are shown in Figure 2B. As expected, measurements returned *α* ≈1 for a bead in water(30), whereas the motion of the tracer was sub-diffusive within embryonic tissues (with *α* = 0.77 for the experiment depicted in Figure 2B)(8, 34).

To determine the optimal duration of a measurement maximizing the signal to noise ratio, we performed an Allan Variance analysis of the equipment, which returned an optimal measurement duration of 2-3 s (38). In aqueous samples laser trapping has been reported heat up the site of the trapped particle by about 1°C/100 mW (39, 40). As our measurements take place deep into highly scattering tissue, the laser intensity reaching the tracer will be significantly less than 300 mW, hence, the temperature increase will be less than 2-3°C. This, in combination with the fact that each measurement lasts 2-3 seconds, renders it likely that temperature effects are negligible.

A classical way of characterizing and comparing viscoelastic properties of different materials independent of the micro-rheological technique employed for the measurement, is by calculating the complex shear modulus, *G*(*f*) (29, 41–44). The complex shear modulus is the ratio of the shear stress to the shear strain, describing the viscoelastic response of a system to a time-dependent stress. It is defined as

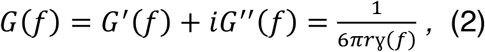

where *r* is the radius of the bead, *G′*(*f*) denotes the storage modulus and describes the elastic response of the system, and *G″*(*f*) denotes the loss modulus and describes the viscous response of the system. *γ*(*f*) is the complex response function and more information on this and on the details of calculating *G* (*f*) are given in the Materials and Methods section. Importantly, the storage and loss moduli scale with frequency by the same exponent describing the scaling of the power spectral data: *G′* ~ *f^α^* and *G″ ~ f^a^* (29). While calculating the complex moduli one should be aware of the inherent frequency limitation originating from the finite maximum measurement frequency employed (detailed in Materials and Methods in the Supporting Information).

As a control, we first calculated the complex shear modulus, *G*(*f*), for a nanoparticle optically trapped in water. The results for a nanoparticle trapped in water (black lines in Figure 2D and supporting Figure S2) show the expected behavior of both the loss and storage moduli in the relevant frequency intervals; in particular, *G″*(*f*) scales with *a*=1 as expected for a purely viscous media where *G* = *G″* = 2*πηf*. To test the feasibility of the method for characterizing cellular viscoelastic properties of internal tissues, we calculated the scaling exponents, α, as well as the storage, *G′*(*f*), and loss, *G″*(*f*), moduli for a subset of gut and liver progenitors (n=5). Both cell populations have a common endodermal origin and are located about 90-120 μm within zebrafish embryos at 30 hpf. Power spectra obtained from liver and gut progenitors scale with frequency in the interval 400 Hz < *f* < 4000 Hz with distinct scaling exponents (*a_gut_* = 0.56±0.08 and *a*_*live*r_=0.76±0.06 for the depicted experiments), thus inferring differential viscoelastic properties for the liver and gut (Figure 3A). Calculation of the complex shear modulus within the same frequency window and of the same type of data as analyzed by power spectral analysis shows that the loss modulus, *G″*(*f*), scales with frequency by exponents of 0.52±0.09 and 0.76±0.05 for gut and liver progenitors, respectively (Figure 2D). As expected, these *a*-values are consistent with those obtained by power spectral analysis. Moreover, trapping beads at 20 and 100 μm depth in Matrigel, representing a material with uniform mechanical properties, returned consistent α-values at both depths, demonstrating that the *a-*value does not depend on the depth of the NIR laser trapped beads (Figure S3). Altogether, these data demonstrate that viscoelastic cell properties in deep embryonic tissues can be quantified *in vivo* based on thermal fluctuation of optically trapped nanoparticles.

**Figure 3.**
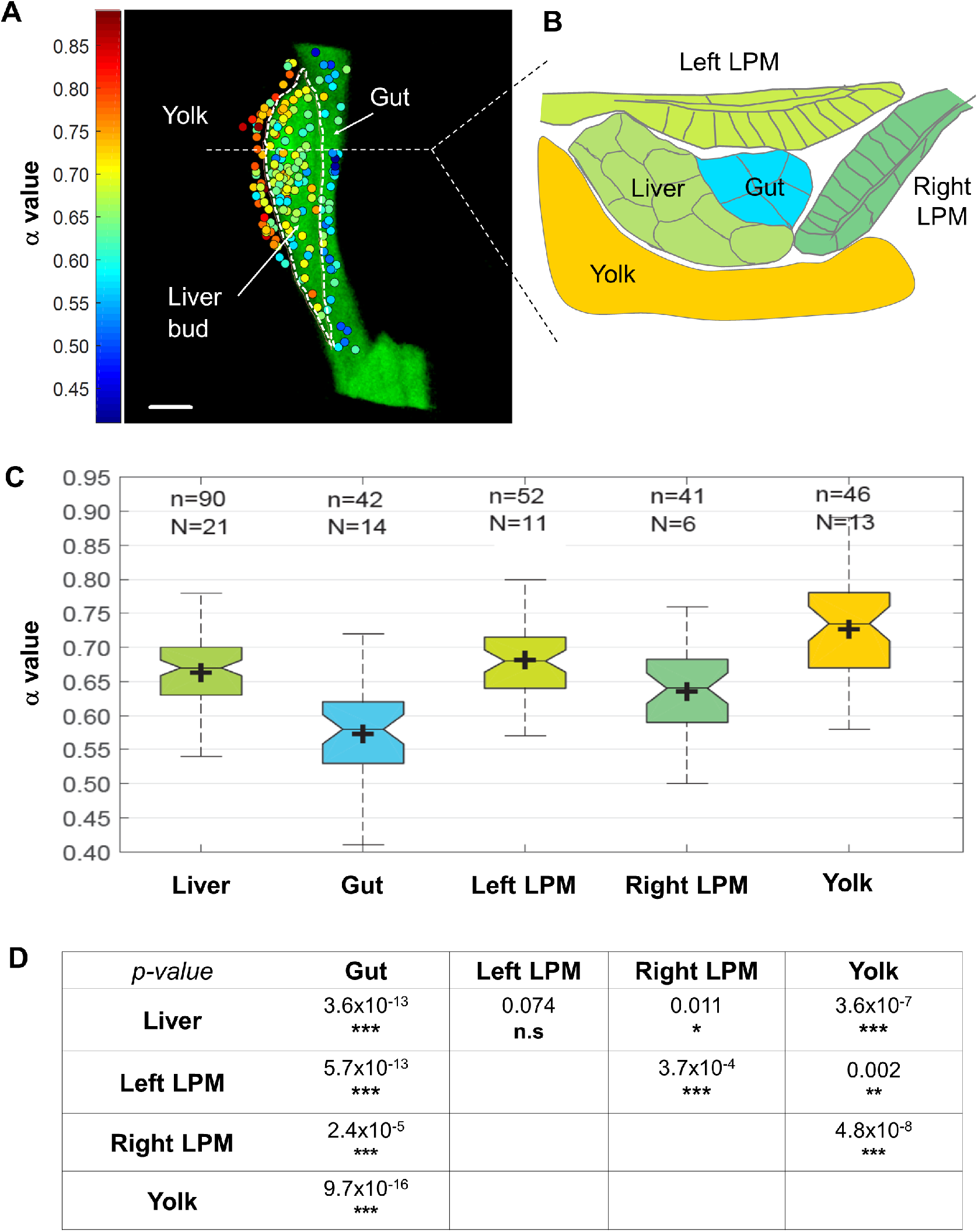
Viscoelastic map of the foregut region reveals different biomechanical properties of neighbouring progenitor populations. (A) 2D projection of viscoelasticity map of the liver, gut, and yolk. The color scale on the left indicates cellular viscoelasticity: increasing blue values show more elastic tissues, while increasing red values represent more viscous tissues (n=178 nanoparticles, N=32 embryos). Scale bar: 20 *μ*m. (B) Viscoelasticity map of the foregut region in a projection orthogonal to (A) and using color code corresponding to average *α* values for each tissue type, color scale bar as in (A). (C) Quantification of *α* values for different tissues shown in a box plot; n = number of analyzed particles, N = number of embryos. (D) Statistical comparison of *α* value distributions between different tissues. The table provides the p-value calculated for each pair of tissues using a two-tailed equal variance Student’s T-test. *p< 0.05, **p<0.01, ***p<0.001

### Cell populations within the foregut region display distinct viscoelastic properties

To quantify the viscoelastic properties of cell populations relevant for foregut morphogenesis, we laser-tracked thermal fluctuations of nanoparticles in the cytoplasm of cells within the developing gut, liver, yolk, or left and right lateral plates inside living zebrafish embryos. After the *in vivo* measurement, the zebrafish embryo was fixed and imaged by confocal microscopy (representative images are shown in Figure 1F and Supporting Figure S4). In the vast majority of the experiments (quantification of all data is shown in Figure S5, one example is shown in Figure S4G-G”), the tracer particle was located in the cytoplasm at a distance away from the nucleus and away from actin cortex that was significantly larger than the amplitude of its typical thermal fluctuation (50 nm).

By coupling the micro-rheological measurement of each nanoparticle to its location within the specific cell population, using the scaling exponent *a* as a measure, we generated a spatially resolved viscoelasticity map of the foregut region. This revealed significant differences between both liver and gut progenitor populations and niche tissues directly adjacent to the liver, including parts of the left and right LPM and the yolk (Figures 3, S6). The cell type-specific *α*-values between individual embryos are highly consistent, corroborating the robustness of the approach (Figure S6).

Despite the common origin from the foregut endoderm, gut progenitors stand out as significantly more elastic compared to liver progenitors with average *a* values of 0.57 (n=42) and 0.66 (n=90), respectively (Figure 3C). Both in the liver and gut regions, the loss modulus dominates the storage modulus (*G″* > *G′*) for all frequencies (Figure 2D), hence, overall the cells in the foregut region have a predominantly liquid-like character. This is supported by the damping factor, 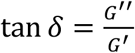 (Figure S7) *G* which for all *f* is higher for the liver than the gut. This, together with the significant difference in the *a*-values between the two populations, demonstrates that progenitors in the liver bud are more viscous than gut progenitors, and gut progenitors are more elastic than liver progenitors.

In the surrounding niche, the adjacent LPM epithelia exhibit similar viscoelasticity as liver progenitors, while the yolk is more viscous than any of the other tissues (Figure 3C-D). Moreover, we identified significantly different *α*-values for the left and right LPM, which share the same embryonic origin and epithelial organization, though move asymmetrically during liver morphogenesis (Figure 3, 1C).

### Microtubule concentrations are higher in gut than liver progenitors

As the cytoplasmic viscoelastic properties of gut progenitors were found to differ significantly from those of liver progenitors in a frequency interval relevant for cytoskeletal biopolymer dynamics, we next investigated which biopolymers could be responsible for this observation. We analysed the distribution of actin filaments and microtubules, as they are major contributors to cytoskeletal stiffness and dynamics. For both tissues, we found that actin localizes mainly to the cell cortex, while the microtubule network is distributed throughout the cytoplasm excluding the volume of the cell nucleus (Figure 4A-C). Notably, microinjected nanoparticles are surrounded by the microtubule network but typically do not co-localize with actin filaments (Figure 4B,C; Figure S4). For both liver and gut progenitors, the majority of nanoparticles are further away from the actin cortex and from the nucleus than the typical amplitude of their thermal fluctuations within the optical trap (~50 nm; Figure S5). This makes the microtubule network the prime candidate for causing differences in viscoelastic properties between progenitor populations on the molecular level.

**Figure 4.**
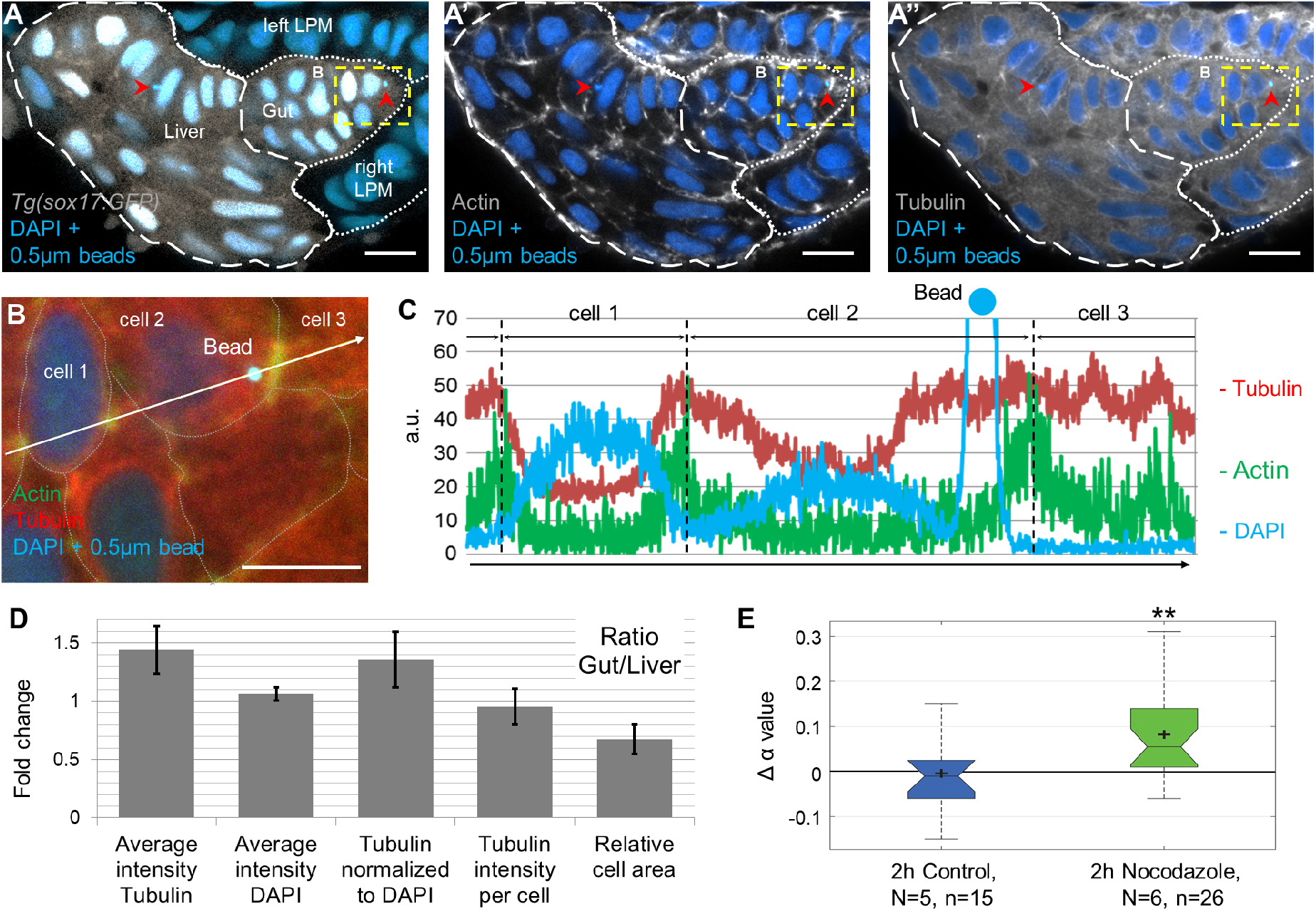
Microtubules surround injected nanoparticles and their concentration is increased in gut progenitors. (A-A”) Transverse section of a 30 hpf *tg(sox17:GFP)* embryo microinjected with 0.5 *μ*m fluorescent polystyrene beads (light blue dots at red arrowheads) showing Phalloidin staining of actin filaments (grey in A’) and β-Tubulin staining to visualize microtubules (grey in A”). Fluorescent emission of microinjected beads and nuclear DAPI occurs at similar wavelengths; beads are detected by size and high signal intensity. Scale bar: 10 *μ*m. (B) Magnification of cellular location of a bead (yellow dashed rectangle in A-A”). Cell borders are outlined based on presence of cortical actin (dashed lines). Scale bar: 20 *μ*m. (C) Intensity profile of Actin, Tubulin and DAPI along the white arrow in B. The high peak in the DAPI channel indicates bead position. (D) Ratio of tubulin levels between gut and liver determined from tissue volumes with a typical linear dimension of 20 *μ*m (see Methods and Figure S5). The average intensity represents the signal intensity normalized to tissue area (tubulin) or nuclear area (DAPI) for each optical section. Tubulin intensity per cell is calculated by normalizing the average tubulin intensity to the amount of nuclei. The relative cell area was obtained by normalizing volume of the respective tissue to nuclei number. Error bars represent one standard deviation, number of analysed 20 *μ*m tissue volumes obtained from different embryos = 4. (E) α-value changes after 2 hours of 2*μ*M Nocodazole treatment (green; mean = 0.082±0.098) and after 2 hours without treatment (blue; mean = −0.004±0.0852). N=number of embryos, n=number of measured nanoparticles; p-value calculated using unpaired equal variance 2-tailed t-test; **p =0.0086.

High cell density and complex 3D arrangement of cells within the foregut region make it difficult to analyse the microtubule cytoskeleton on the single cell level. Therefore, we quantified average microtubule density for gut and liver progenitors on the tissue level. To avoid region-specific and cell orientation bias, analyses were performed on volumes with linear dimensions of 20 *μ*m, yielding cumulative values from individual optical sections. The relative amount of microtubule is 1.44 +0.21 - fold higher in gut compared to liver progenitors (Figure 4D; Figure S4E). Intensity of nuclear DAPI of the respective tissues was used to correct for a potential bias during data acquisition. This revealed that the amount of microtubule per cell is similar between the two populations (0.96 +0.06 - fold). However, the area of gut progenitors is only 0.67 +0.24 - fold of that of liver progenitors. Hence, despite a similar cellular amount of microtubules in the two populations, the microtubule concentration within gut progenitors is about 1.36 +0.24-fold higher per cell.

To assess whether the microtubule cytoskeleton influences viscoelastic cell properties we disrupted microtubule polymerization using drug treatments with 2*μ*M Nocodazole. We measured *α*-values for the same nanoparticles located in gut progenitors before (at 28 hpf) and after 2 hours of drug treatment. In controls, particles were measured at the same 2-hour interval without drug administration. Microtubule destabilization lead to a significant increase in the *α*-value indicating a shift to more viscous cell properties (Figure 4E, Figure S8A,B). Concomitantly, microtubule destabilization during foregut morphogenesis impairs asymmetric gut looping and liver bud formation (Figure S8C-F). Thus, the microtubule cytoskeleton influences viscoelastic cell properties and microtubule concentration correlates with the differential elasticity between gut and liver progenitors.

## DISCUSSION

We performed nearly non-invasive quantitative micro-rheological investigation of cells and tissues in a living embryo at unprecedented depth. Using the cell populations within the developing foregut region as a model to investigate material properties *in vivo*, we show that cells exhibit consistent viscoelasticity within a population, whereas viscoelasticity significantly differs between various cell types, with gut progenitors being more elastic than any other population in the region. On the molecular level, we show that the more elastic property of gut progenitors correlates with a higher concentration of microtubules in comparison to the liver progenitors and that microtubule disruption alters cellular viscoelasticity, as well as foregut morphogenesis.

In the laser-tracking experiments, we used a sampling frequency of 22 kHz, as this is the frequency minimizing the Allan Variance of our setup (38). However, the quadrant photodiode used for data acquisition can sample with frequencies up to 100 MHz. Hence, even without pushing the limits of the quadrant photodiode, this type of data acquisition is several orders of magnitude faster than previously published particle tracking methods based on video recording (15, 18, 21, 45, 46), thereby allowing for a significant expansion of the timescale from which information can be retrieved.

The intracellular tracer nanoparticles used in this study are found not to be directly adjacent neither to the cell cortex nor to the nucleus, thus, they mainly provide information about the viscoelastic properties of the cytoplasm. For both liver and gut we find *G″ > G′* at all frequencies, hence, for the intracellular environment in the developing gut region the viscous properties dominate over the elastic. Our data for cells located in internal tissues 90-150 *μ*m deep within zebrafish embryos agree with video-based tracer measurements inside the *Drosophila* syncytium (21) which were performed at depths up to 40 mm in the frequency interval 0<*f*<1000 Hz. In accordance with our findings, the cytoplasmic viscoelastic properties in *Drosophila* embryos were shown to be dependent on microtubules rather than the actin cytoskeleton. In contrast, atomic force microscopy (AFM)-based microrheology performed on isolated cells report *G′ > G″* for similar frequencies (47–49). This is likely due to the extracellular location of the probe as opposed to our method; an AFM operates on the cell surface and cortical actomyosin thereby significantly contributes to such micro-rheological measurements (Brückner et al., 2017; Rigato et al., 2017).

AFM and techniques such as microfluidics-based assays (50) or optical stretching (51) can measure viscoelasticity of isolated cells and tissues, however, they cannot be directly used for internal tissues *in vivo*. Other methods described to investigate biomechanical properties in deep tissues, including Brillouin microscopy or tomography (52, 53),report primarily on the length scale of tissues. In contrast, the presented laser-based assay causes no detectable physiological damage and can be used to probe single cells in intact tissues and embryos without disrupting their native environment. By laser-tracking intracellular tracer-nanoparticles, we demonstrate that different cell populations have distinct viscoelastic properties indicating that cytoplasmic viscoelasticity is cell-type specific and can be used to distinguish cell populations *in vivo*.

The optically tracked nanoparticle returns information about the local viscoelastic environment inside living cells. The application of this technique could be straightforwardly expanded to examine tracers located in the extracellular space, for instance in the extracellular matrix, which is of interest due to its importance for differentiation of embryonic and induced stem cells (54). In addition, this method can be adapted for active micro-rheological and force measurements (19), as optical trapping allows manipulation of the particle location. Due to the high penetration of NIR light, the method should be applicable to any organism or tissue amenable to NIR, and therefore should be suitable for probing biomechanical properties *in vivo* in species other than zebrafish, as well as complex *in vitro* cultures, such as explants and organoids.

The timescales here investigated are those relevant for cytoskeletal dynamics, including microtubule turnover and polymerization. Recent work shows cell autonomous functions for microtubule-mediated mechanics in the developing *Drosophila* wing epithelium, thus providing evidence for the contribution of material properties to tissue scale morphogenesis (55). At the stage of liver bud formation, gut progenitors exhibit little to no motility and shortly after give rise to the intestinal epithelium. At the same time, however, both liver progenitors and LPM undergo dramatic cellular rearrangement before differentiating into their respective cell types. The comparatively fluid nature of the liver progenitors and LPM epithelia corroborates their active migration and movement (23, 24). In contrast, higher tubulin concentration of the more static gut progenitors, as found in the current study, is consistent with an increased stiffness to counteract the dynamic rearrangement of surrounding tissues, given that microtubules are the stiffest cytoskeletal filaments (56). Consistently, we find that microtubule disruption results in decreased gut progenitor elasticity and altered foregut morphology. Furthermore, our results in conjunction with lower cell rearrangement among gut than liver progenitors (23), are comparable to cell behaviours associated with tissue stiffness during axis elongation in zebrafish (57). A similar relationship between motility and viscoelasticity has also been observed in cancer spheroids, where the motile cells located at the invasive tips of the spheroid are more viscous than those statically located at the base which are more elastic (2). In our work we find the yolk adjacent to the forming foregut to be the most viscous region probed, indicating an environment that is permissive to dynamic tissue rearrangements. Altogether these results suggest that differences in material properties between adjacent developing tissues may drive or facilitate cell movement and/or tissue differentiation.

In summary, we present a method for quantifying and mapping material properties in cells and tissues *in vivo*, which is a prerequisite for establishing a firm connection between material properties, biomechanics and cell behaviors in development and disease.

## METHODS

### Experimental Model and Subject Details

Adult zebrafish and embryos were raised according to standard laboratory conditions(58), and all experiments were performed in agreement with the ethical guidelines approved by the Danish Animal Experiments Inspectorate (Dyreforsøgstilsynet). The transgenic line Tg(−0.5 sox17:GFP)zf99 was used to visualize the endoderm(59). To prevent pigment formation, after 24 hours post fertilization (hpf) embryo medium (120 mg/L sea salt; Instant Ocean, Aquarium Systems, France) was supplemented with 0.2 mM PTU (1-phenyl-2-thiourea; Sigma-Aldrich, USA).

### Preparation of zebrafish embryos

0.5 *μ*m polystyrene fluorescent beads (FP-0545-2; SPHERO Fluorescent Particles, Light Yellow, 0.4-0.6 *μ*m; Spherotech, USA) were injected at the 1-cell stage into the cytoplasm-yolk interface. Prior to injections, the stock solution of fluorescent particles (1% w/v in deionized water with 0.01% NP40 and 0.02% Sodium Azide) was diluted 1:10 in autoclaved deionized water, and 0.5-1 nl was injected per embryo to ensure a sufficient number of beads in the foregut region at 28-30 hpf and reduced extracellular clusters. At 28-30 hpf, injected embryos were manually dechorionated and anesthetized by incubation with 0.14 mg/ml Tricaine (Ethyl 3-aminobenzoate methanesulfonate; A5040, Sigma-Aldrich, USA).

Embryos were embedded dorsolaterally on the left side, with the liver bud oriented towards the coverslip (25×60 mm #1 coverslip, Menzel-Gläser, Germany), in a drop of 0.4% low melting temperature agarose (NuSieve GTG Agarose; Lonza, USA).

Before agarose polymerization, embedded embryos were covered with another coverslip (24×40 mm #1; Menzel-Gläser, Germany). The two coverslips were separated by vacuum grease (Dow Corning High Vacuum Grease) applied on the longitudinal edges of the second coverslip. After agarose polymerization, the remaining space in the chamber between the coverslips was filled with approximately 200 *μ*l embryo medium containing PTU and Tricaine, as described above. Finally, the chamber was sealed by vacuum grease.

### Optical trapping and confocal imaging

A near infrared laser beam (λ=1064 nm, Nd:YVO4, Spectra-Physics J20-BL106Q) was implemented into a Leica SP5 confocal microscope (Figure 2A), thus allowing for simultaneous optical trapping and visualization; for details of the setup see reference (28). A Leica (PL APO, NA = 1.2, 63X) water immersion objective was used for focusing the trapping laser beam inside the sample as well as for confocal image acquisition. Forward-scattered laser light was collected by an immersion oil condenser, located above the sample, and imaged onto a Si-PIN quadrant photodiode (Hamamatsu S5981) placed in the back focal plane. Data were collected using custom-made Labview programs using a sampling frequency of 22 kHz.

In parallel with optical trapping of the nanoparticles, confocal imaging was performed using excitation lasers with wavelengths of 405 nm and 488 nm, thereby exciting fluorophores on the nanoparticles and visualizing the foregut endoderm. For all measurements, the laser was operated for 2-3 s, thus minimizing the Allan Variance of the setup(38) and thereby maximizing the signal to noise ratio, and using power of ~300 mW in the sample. In agreement with the literature(13), no physiological damage was observed as a consequence of this irradiation. Also, the expected temperature increase during the 2-3 s measurement interval is well below 2-3 °C(40). All measurements were performed at room temperature.

During measurements, we visually in three dimensions co-localized the focus of the optical trap with a single nanoparticle inside the fish. Importantly, the amplitude of the power spectrum significantly increased upon nanoparticle trapping compared to having an empty trap inside the fish (Supporting Figure S1), because upon correct alignment of the optical trapping system, a nanoparticle in the focus of the trap serves as a lens increasing the number of photons reaching the QPD. Using a 3D piezo-stage (Mad City Labs) the position of the optical trap was fine-tuned in order to maximize the amplitude of the power spectrum and the measurement was acquired with these settings.

### Localization of injected nanoparticles

To analyze overall nanoparticle distribution, cell membranes were mosaically labelled by co-injecting Tol2-Ubi-mKate-CAAX plasmid DNA (courtesy of Sara Caviglia). Liver progenitors were visualized by the immunostaining for EfnB1(23). To determine precise location of nanoparticles used for measuring viscoelasticity, the embryos were fixed in 4% paraformaldehyde (PFA) immediately after the optical trapping experiments, deyolked and stained with phalloidin conjugated to Atto 633 (Sigma-Aldrich, USA) to visualize the actin cell cortex and tissue morphology. The foregut region was imaged using a Leica SP8 confocal microscope. Images obtained from confocal scanning simultaneously with the trapping experiments were used to identify the particles of interest in the fixed sample by manually correlating foregut shape and nanoparticle distribution in 3D using Imaris BITPLANE software. The nanoparticles were assigned to a specific cell population based on tissue morphology and transgenic sox17:GFP expression. Only single beads, which could be clearly assigned to a specific tissue, were included in the analysis.

### Visualization of microtubules and actin filaments

Immunostaining for β-tubulin was used to visualize microtubules, while actin filaments were stained using Phalloidin-Atto 633 fluorophore (Sigma-Aldrich, St. Louis, USA). Immunostaining was performed on sections to reduce penetration bias. Embryos were fixed at 30 hpf in 4% paraformaldehyde (PFA) overnight at 4°C, washed in 1x PBS (Gibco, Life technologies), deyolked and embedded in 4% agarose (Ultra Pure Agarose, Invitrogen) for subsequent sectioning. Transverse 50 *μ*m sections were obtained using Leica microtome and stained using mouse monoclonal β-tubulin antibody (E7, DSHB; 1-2 *μ*g/ml) followed by secondary goat anti-mouse Cy3 antibody (Jackson ImmunoResearch; 1:300). To visualize actin filaments in embryos microinjected with 0.5 *μ*m nanoparticles, sections were costained with phalloidin-Atto 633 (1:300) and DAPI (1:1000). To discriminate gut and liver for quantification of microtubule density, embryos were co-stained with either rabbit polyclonal EfnB1 (Cayuso et al., 2016; 1:1000) or Prox1 (AngioBio) antibody, followed by goat anti-rabbit Alexa 633 (Jackson ImmunoResearch; 1:300). For confocal imaging sections were mounted in VECTASHIELD Antifade Mounting Medium (Vector Laboratories, USA) on the glass slides (Superfrost Plus; Menzel-Gläser, Germany) and covered with #1 coverslip (Menzel-Gläser, Germany). Consecutive optical sections were collected every 0.33 μm using a Leica SP8 confocal microscope.

### Drug treatment for microtubule depolymerization

Embryos were embedded as described above and nanoparticles in gut progenitors were used for measurements at 28 hpf. After the measurements, incubation medium in the chamber was replaced with about 200 *μ*l embryo medium containing 2*μ*M Nocodazole (Sigma-Aldrich, USA), 0.2 mM PTU and 0.14 mg/ml Tricaine. After 2 hours of incubation at room temperature (average 24°C), measurements were performed on the same nanoparticles. Embryos embedded in agarose remained in the same position during the medium exchange, which facilitated localization of previously measured nanoparticles. For controls, the same nanoparticles were used for measurements at 28 hpf and 2 hours later without Nocodazole incubation.

### Probing viscoelasticity in depth in a uniform viscoelastic material

1 mm beads were optically trapped at different depths inside a DMEM: Matrigel (in a ratio of 3:1) viscoelastic matrix. To prepare the matrix, frozen matrigel was slowly thawed on ice and mixed with cooled down DMEM and nanoparticle-solution. It was injected between two coverslips and the chamber was sealed with vacuum grease. The chamber was placed on the microscope stage and microbeads were trapped at depths of 20 and 100 μm inside the matrix. The obtained measurements were analyzed as described below and the data are shown in supporting Figure S3.

## ACKNOWLEDGMENTS

We thank Charlotte Bailey, Sara Caviglia, Jakub Sedzinski and Raghavan Thiagarajan for their helpful comments on the manuscript and S. C. for *ubi:CAAX-mKate* plasmid. This work is supported by the Danish National Research Foundation via the StemPhys CoE (DNRF116) and by the Novo Nordisk Foundation to E.A.O. (NNF17CC0027852).

## Notes

### Competing Interest Statement

The authors have declared no competing interest.

