## Supplemental Information for "Foregut organ progenitors and their niche display distinct viscoelastic properties *in vivo* during early morphogenesis stages"

### METHODS

#### Data quantification and statistical analysis

<u>Power spectral analysis and calculation of the complex shear modulus</u>

The QPD yields a position measure in arbitrary units (Volts). To obtain absolute values of the complex shear modulus it is necessary to convert these arbitrary units to SI units (meters). To do this, we determined the conversion factor, *b*, relating measurements in Volts to measurements of distance, *x*, in meters: $x = \beta V_x$. *b* can be extracted from the power spectrum of a trapped bead in water:

$$P(f) = \frac{K_B T}{6\pi\eta r\beta^2({f_c}^2+f^2)} \quad , \qquad (1)$$

where *T* is the temperature, $K_B$ is the Boltzmann constant, $\eta$ is the viscosity of the medium, *r* is the radius of the particle, and $f_c$ is the corner frequency, *$f_c$=k/2pg*, where *k* is the trap stiffness, characterizing the harmonic potential of the optical trap, and *g* is the friction coefficient.

Inside a viscoelastic environment the scaling of the power spectra scale with frequency, *f,* is characterized by the scaling exponent *a*: $P_x(f) \equiv \langle|\tilde{x}(f)|^2\rangle \propto f^{-(1+\alpha)}$ (where, trivially, *a*= 1 for a tracer in water). All power spectra were calculated by custom written MATLAB programs. An average of 3 measurements in *x* and *y* directions was used to extract the value of *α* for each trapped nanoparticle in a single cell. The average of a values obtained for a specific cell type was used to infer the viscoelastic properties of the corresponding tissue. Statistical significance for differences in viscoelasticity between tissue pairs was determined using a two-tailed equal variance Student's T-test. The significance level was set at 5%. The number of embryos (N) and sample size (n) are indicated in the legend to Figure 5.

The sample size was not computed and decided at the stage of the study design, as the levels of variation and the difference between the samples were hard to predict. However, based on previous publications that used other techniques to measure cell viscoelasticity, we estimated N≥5, n≥40 for unpaired samples (different cell populations) and n≥15 for paired samples (drug treatment) would be reasonable numbers for sample sizes.

To calculate the complex shear modulus, $G(f)$, we note that the Fourier transform of the position of a bead in a viscoelastic medium, $x(f)$, is linearly related to the Fourier transform of the thermal force, $F(f)$ (1):

$$x(f) = \gamma(f)F(f) \ . \qquad (2)$$

Here, $\gamma(f) = \gamma'(f) + i\gamma''(f)$ is the complex response function from which the complex shear modulus, $G(f)$, can be found through the Stokes–Einstein relation:

$$G(f) = G'(f) + iG''(f) = \frac{1}{6\pi r \gamma(f)} \qquad (3)$$

where $r$ is the radius of the bead, $G'(f)$ denotes the storage modulus, and $G''(f)$ denotes the loss modulus. For a bead in an optical trap, $G'(f)$ is the sum of the storage modulus of the medium and of the optical trap, which exerts a harmonic potential on the particle. If the trap is sufficiently weak, like in current experimental conditions, the elastic contribution from the trap can be ignored.

The imaginary part of the response function can be extracted from power spectral density of the trapped bead, $P(f)$:

$$\gamma''(f) = \frac{\pi f}{2K_B T} P(f)\ . \qquad (4)$$

From this, the real part of $\gamma(f)$ can be calculated using a Kramers–Kronig transformation:

$$\gamma'(f) = \frac{2}{\pi} P \int_{f''=0}^{f''=\infty} df'' \frac{f''\gamma''(f'')}{f''^2 - f^2}, \qquad (5)$$

where *P* stands for the principal value of the integral. This integral can be solved numerically, thus retrieving $G(f)$.
Both storage and loss moduli scale with frequency by $\alpha$ , the same scaling exponent describing the scaling of the power spectral data: $G' \sim f^{\alpha}$ and $G'' \sim f^{\alpha}$ (2) . When analyzing real experimental measurements, an inherent experimental frequency limitation, due to the existence of a finite maximum measurement frequency, causes errors to occur in $\gamma'(f)$ at sufficiently high frequencies. For the shear modulus, such errors have been shown to be negligible at frequencies below $f_{Nyquist}/2$, where $f_{Nyquist}$ is the Nyquist frequency. For the storage modulus, the error is only negligible for frequencies $f \leq f_{Nyquist}/10$ (3). In our experiments, the QPD was operated at 22 kHz, hence, $f_{Nyquist}$ = 11 kHz.
Using equations (4) and (5), we determined the imaginary and real part of the response function for 10 < *f* < 10 kHz. Via equation (3) the loss and storage moduli were calculated.

Quantification of microtubule distribution

60 consecutive optical sections obtained every 0.33 $\mu$m were used to quantify relative microtubule density in gut and liver for a single embryo. Tissues were manually outlined in Fiji (4) based on *Tg(sox17:GFP)* expression in the endoderm and staining for EfnB1 or Prox1 as a marker for liver progenitors. For higher efficiency, outlining was performed on 1 $\mu$m cumulative projections obtained by combining pixel intensity of 3 consecutive 0.33 $\mu$m optical sections (in Fiji: Image->Stacks->Z-projection->Sum Slices). Thus, for a single embryo 20 cumulative projections were obtained from 60 consecutive 0.33 $\mu$m optical sections corresponding to 20 $\mu$m volume. Nuclei were outlined by creating a binary mask based on DAPI intensity with the same threshold for all cumulative projections within a 20 $\mu$m volume. Intensity and area were quantified by plotting a histogram of the

outlined regions for each cumulative projection (macro in Fiji: run ("Histogram", "bins=1000 x_min=0 x_max=1000 y_max=Auto")), which returns the number of pixels with a specific value of signal intensity. The bin size was set to 1000, as cumulative projections were generated as a sum of three 8 bit images, and the intensity of a pixel should be ≤ 3x255=765. The area was calculated as a sum of all pixels within the outlined region, while the total signal intensity was obtained as a sum of signal intensity values multiplied by the corresponding amount of pixels. The total signal intensity and area within a 20 $\mu$m volume for gut and liver, as well as for the corresponding nuclei, were quantified as sums of individual cumulative projections. The average signal intensity represents the total signal intensity divided by the area. “Tubulin normalized to DAPI” in Figure 6 represents the average tubulin intensity divided by the average DAPI intensity. Tubulin intensity per cell was calculated as the average tubulin intensity divided by the amount of nuclei manually calculated within the 20 $\mu$m volume. The relative cell area represents the area calculated for gut or liver divided by the amount of nuclei within the corresponding tissue. The ratio between gut and liver was calculated for each 20 $\mu$m volume individually, and average values represent 4 volumes obtained from different embryos.

### Resources and vendors:

| REAGENT or RESOURCE | SOURCE | IDENTIFIER |
|---|---|---|
| Antibodies | | |
| Polyclonal rabbit anti-EphrinB1 | Cayuso et al.,2016 | |
| Polyclonal rabbit anti-Prox1 | AngioBio | Catalog #: 11-002 |
| Monoclonal mouse anti-β-tubulin | DSHB; (5) | E7 |
| goat anti-mouse Cy3 | Jackson ImmunoResearch Laboratories | 115-166-146 |
| goat anti-rabbit Alexa647 | Jackson ImmunoResearch Laboratories | 111-606-003 |
| Chemicals, Peptides, and Recombinant Proteins | | |
| 1-Phenyl-2-thiourea (PTU) | Sigma-Aldrich | P7629; CAS: 103-85-5 |
| Ethyl 3-aminobenzoate methanesulfonate (Tricaine) | Sigma-Aldrich | A5040; CAS: 886-86-2 |
| Instant Ocean sea salt | Aquarium Systems | |
| SPHERO Fluorescent Particles, Light Yellow, 0.4-0.6 $\mu$m | Spherotech | FP-0545-2 |
| NuSieve GTG Agarose, low melting temperature | Lonza | Catalog #: 50085 |
| Dow Corning High Vacuum Grease | Højstrup Industrilim | DC5940-0050 |

| | | |
|---|---|---|
| Phalloidin–Atto 633 | Sigma-Aldrich | 68825 |
| Ultra Pure Agarose, for embedding | Invitrogen | 16500-500 |
| PBS Tablets | Gibco | 18912-014 |
| VECTASHIELD Antifade Mounting Medium | Vector Laboratories | H-1000 |
| Experimental Models: Organisms/Strains | | |
| *Tg(-0.5 sox17:GFP)$^{zf99}$* | Mizoguchi et al., 2008 | |
| Recombinant DNA | | |
| Tol2-Ubi-mKate-CAAX | Courtesy of Sara Caviglia and E.A.O | |
| Software and Algorithms | | |
| Bitplane IMARIS | Bitplane | http://www.bitplane.com/imarist |
| Labview | National Instruments | http://www.ni.com/labview/release-archive/2010/ |
| LAS AF Lite | Leica | https://www.leica-microsystems.com |
| Matlab 2016b | MATLAB | https://www.mathworks.com |
| Fiji | (4) | http://imagej.net/Fiji |
| Other | | |
| Nd:YVO4 (λ=1064 nm laser) | Spectra Physics | https://www.spectra-physics.com/ |
| TCS SP5 microscope used for live imaging during optical trapping | | https://www.leica-microsystems.com |
| TCS SP8 microscope used for fixed tissue imaging | Leica microsystems | https://www.leica-microsystems.com |
| Piezoelectric stage | Mad City Labs | http://www.madcitylabs.com |
| Photodiode | Hamamatsu | S5981 |
| The Leica VT1000 S microtome | Leica Biosystems | https://www.leicabiosystems.com |
| Zeiss LSM 880 with Airyscan for high resolution fixed tissue imaging | ZEISS | https://www.zeiss.com |

### Supporting Figures

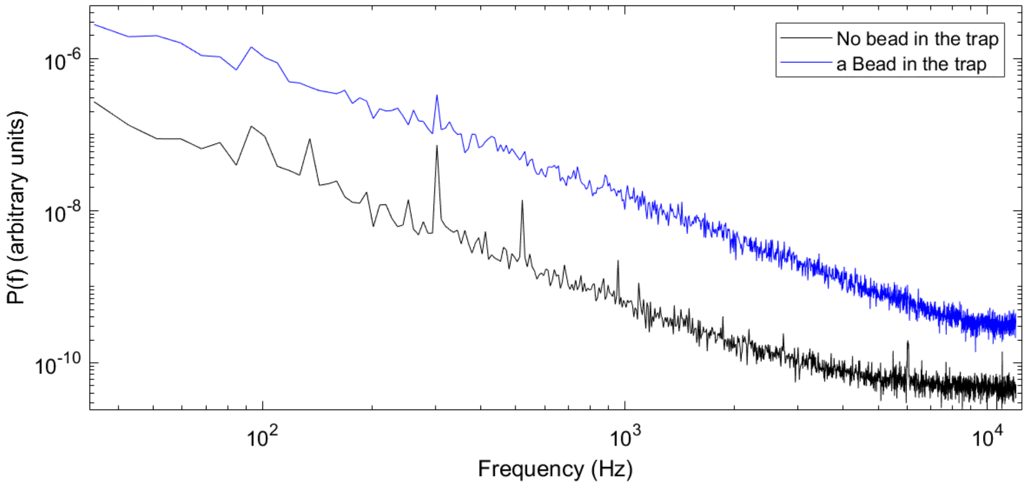

**Figure S1: Power spectra from measurements inside living zebrafish** with (blue) or without (black) a trapped nanoparticle. Due to the focusing effect of the trapped nanoparticle, the amplitude of the power spectrum increases if a nanoparticle is located in the center of the optical trap. The increase in the power spectrum corresponds to an increased signal to noise ratio. Supporting this, noise peaks, e.g., at 300 Hz, are less pronounced upon trapping of a bead.

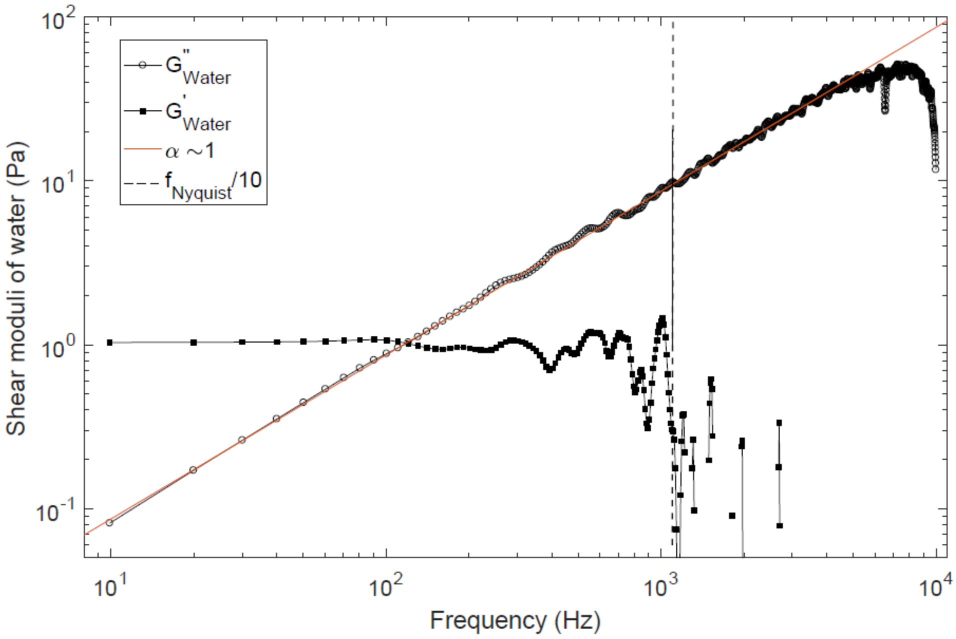

**Figure S2: Complex shear modulus measured by optical trapping of a bead in water.** The harmonic potential of the optical trap gives rise to a storage modulus, *G'* (full squares), that is frequency independent and comparatively small (~1 Pa). As expected for water, a purely viscous liquid, the loss modulus, *G"* (hollow circles), is linearly increasing with frequency with an exponent of *a*=1 (red line).

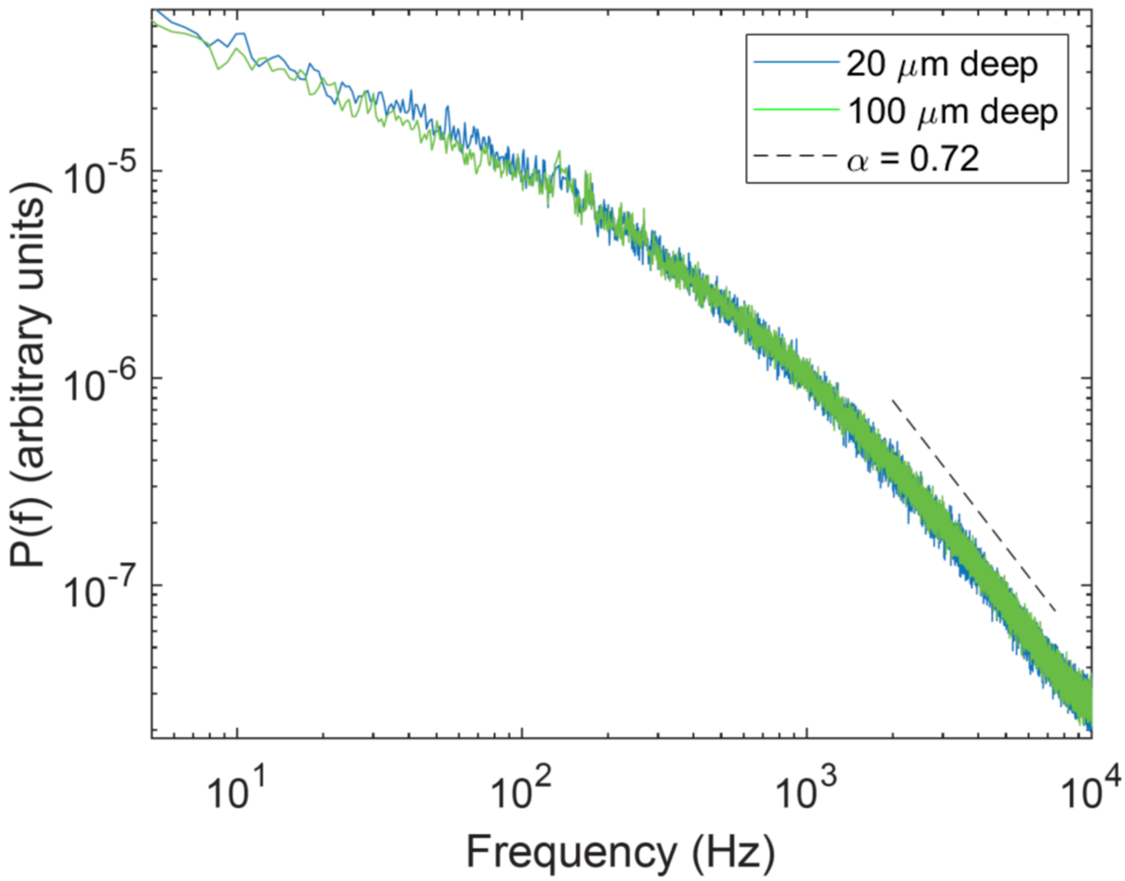

**Figure S3: Scaling exponents, *a*, are independent of trapping depth in a uniform viscoelasatic medium.** Average power spectra for trapped beads at depths of 20 µm (blue) and 100 µm (green) in a uniform viscoelastic matrigel. The slope at relevant frequencies (dashed gray line) is the same at both depths. The data represent an average of 10 experiments for each depth.

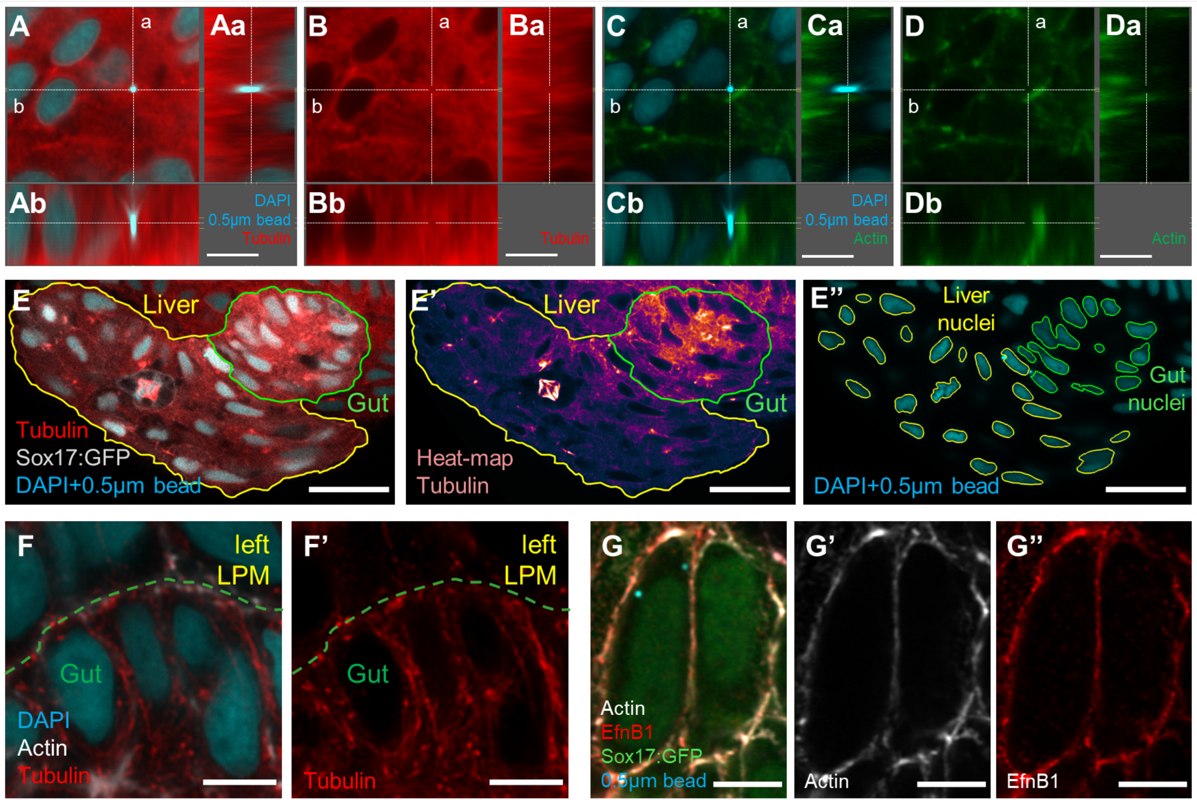

**Figure S4: Analysis of cellular composition by confocal microscopy.** (A-D) Orthogonal projections corresponding to the confocal image shown in Figure 4B. A microinjected nanoparticle (cyan) is surrounded by microtubules (A-B), but not actin filaments (C-D). Scale bar: 5 $\mu$m. (E-E'') Representative 1 $\mu$m summed projection used to quantify tubulin intensity as shown in Figure 4D. (E) Gut (green line) and liver (yellow line) were manually outlined for each 1 $\mu$m summed projection within 20 $\mu$m volumes to quantify corresponding tubulin intensity and tissue area. (E') Signal intensity heat-map demonstrates tubulin enrichment in the gut. (E'') Nuclei were outlined by creating a binary mask based on DAPI staining. Nuclear DAPI intensity (as reported in Figure 4D) for both gut and liver was quantified in the outlined nuclei of gut (green) and liver (yellow) progenitors. Scale bar: 20 $\mu$m. (F, F') Single optical section showing high-resolution views of microtubules obtained with LSM 880 Airyscan. Dashed line marks the border between gut progenitors and left LPM. Scale bar: 5 $\mu$m. (G-G'') Nanoparticles (blue in G) distribute in the cytoplasm between cortical actin and the nucleus as analysed in Supporting Figure S5; single optical section showing cortical actin filaments (grey) close to the plasma membrane marked by EfnB1 (red) in liver progenitors obtained with LSM 880 Airyscan. Nanoparticles (blue) are localized in the cytoplasm. Scale bar: 5 $\mu$m.

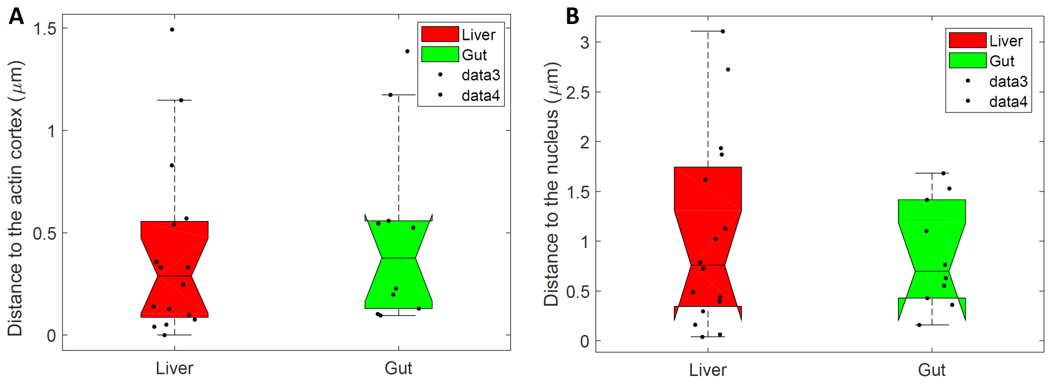

**Figure S5. Location of tracer particles with respect to the nucleus or actin cortex.** High resolution images (see Supporting Figure S4) were used to measure the distance from the tracer particles to the actin cortex (A) or to the nucleus (B) for particles located in liver (red, n=16) or gut (green, n=10) cells, respectively. The vast majority of particles investigated, regardless of cell type, are further away from the nucleus and from the periphery of the cell (actin cortex) than the amplitude of their thermal motion (~50 nm).

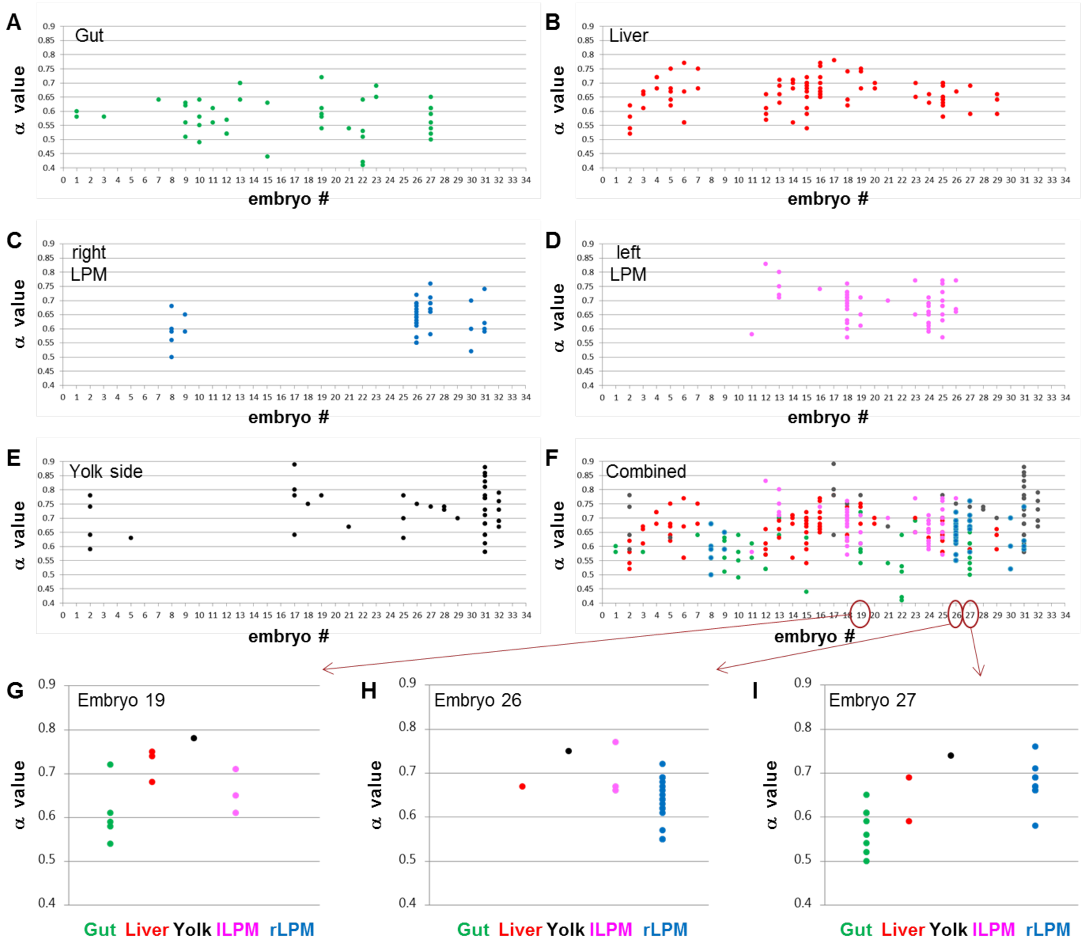

**Figure S6. Relative distribution of $\alpha$ values in different cell populations is consistent between embryos.** Each dot corresponds to the $\alpha$ value measured for a single bead in gut (A, green) or liver (B, red) progenitors, right LPM (C, blue), left LPM (D, magenta), or the yolk (E, black) in one of 32 analysed embryos. (F) Combined distribution of *a* values of all tissues. (G-I) Relative distribution of $\alpha$ values in three representative individual embryos, grouped by tissue type.

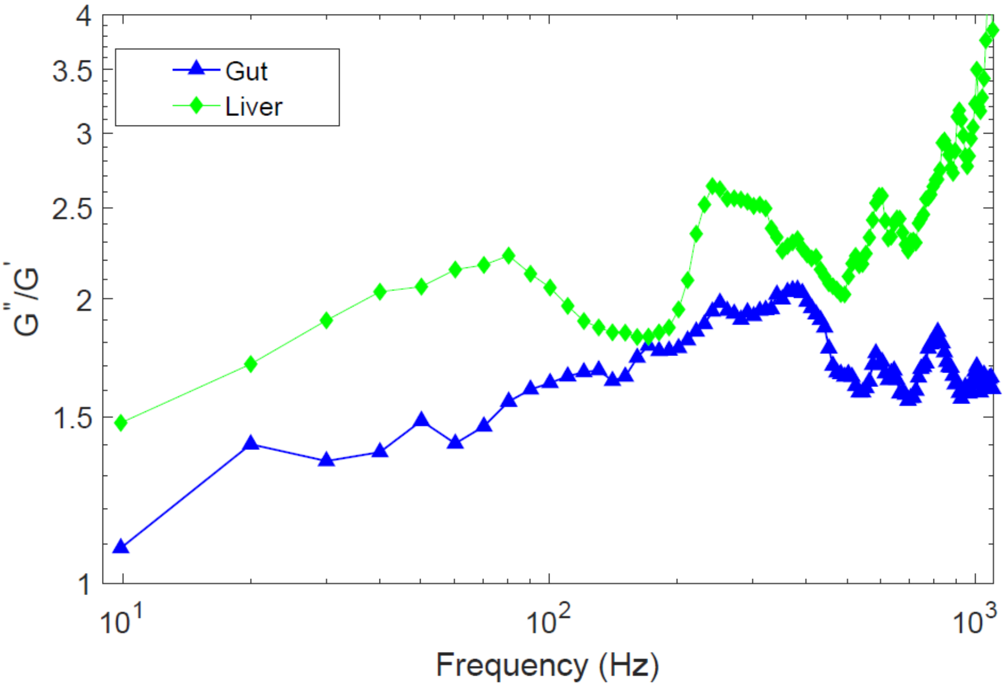

**Figure S7: The damping factor, *G''/G'*, as a function of frequency for the liver and gut area.** At all frequencies, the damping for both the gut (blue triangles) and liver (green diamonds) are larger than 1, thus indicating a predominantly liquid like behavior of these tissues. Also, it is evident that the damping factor overall is larger for liver progenitors, thus indicating their more viscous nature, consistent with the power spectral analysis.

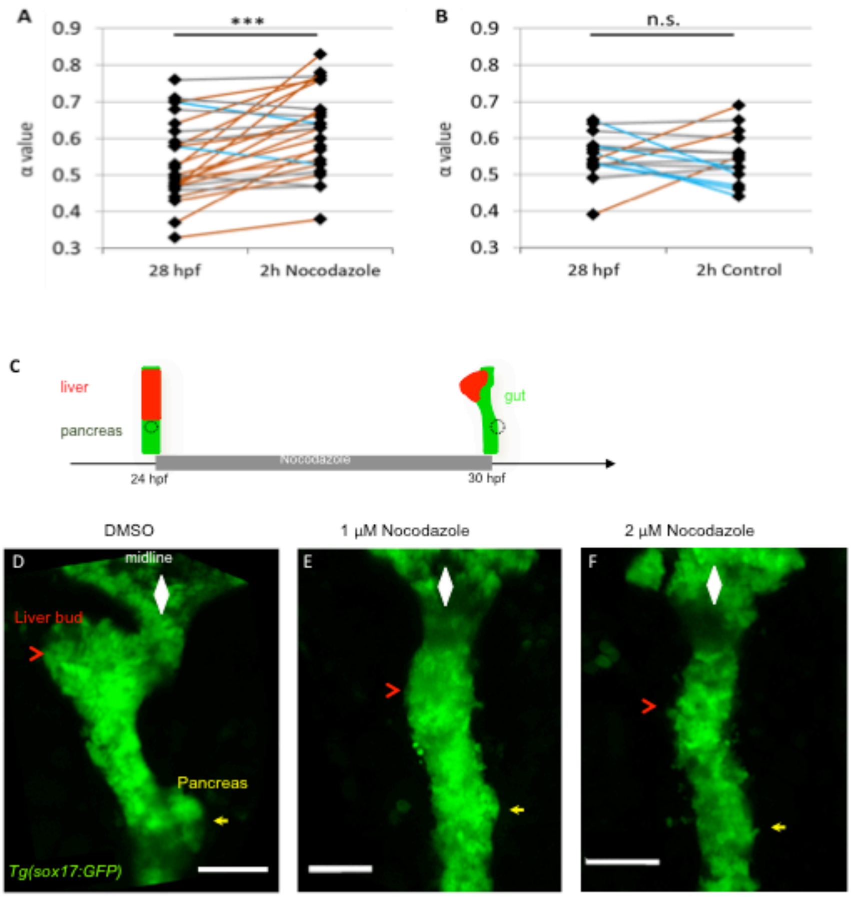

**Figure S8. Disrupting microtubule polymerisation alters cellular viscoelasticity and foregut morphogenesis.** (A,B) α-values determined by sequential measurements of beads in the gut progenitors at 28 hpf and 30 hpf, after 2 hours of 2μM Nocodazole treatment (A, N=6, n=26) or after 2 hours without treatment in controls (B, N=5, n=15). Orange lines indicate α-value changes ≥ 0.05; grey lines indicate α-value change -0.05 < difference < 0.05, blue lines indicate α-value change ≤ -0.05. N=number of embryos, n= number of measured nanoparticles; paired sample 2-tailed t-test: ***p=0.0003, n.s. p=0.8631. (C) Schematic of Nocodazole treatment during gut looping and asymmetric liver bud formation. (D-F) Ventral views of *tg(sox17:GFP)* show how 1 μM and 2 μM Nocodazole treatments impair gut looping and liver bud formation compared to the control (D) undergoing only DMSO treatment; anterior to the top. n=10, N=2.

**Supplementary movie: Optical tweezers trapping a bead in the foregut region of *live* zebrafish.** The position of the trap is indicated by a red circle and red points show the locations of beads in the foregut region. The bead-injected embryo was moved very slowly from the left to the right side of the trap, which was kept at a constant position. When a bead came close to the trap, it was pulled into the trap and remained trapped at a constant position for several seconds while the embryo continued to move.
